# Phospholipids Diffusion on the Surface of Model Lipid Droplets

**DOI:** 10.1101/2022.06.30.498225

**Authors:** Shima Asfia, Ralf Seemann, Jean-Baptiste Fleury

**Author notes:** Corresponding author, *Email address:* (Jean-Baptiste Fleury).

## Abstract

Lipid droplets (LD) are organelles localized in the membrane of the Endoplasmic Reticulum (ER) that play an important role in metabolic functions. They consist of a core of neutral lipids surrounded by a monolayer of phosphoplipids and proteins resembling an oil-in-water emulsion droplet. Many studies have focused on the biophysical properties of these LDs. However, despite numerous efforts, we are lacking information on the mobility of phospholipids on the LDs surface, although they may play a key role in the protein distribution. In this article, we developed a microfluidic setup that allows the formation of a triolein-buffer interface decorated with a phospholipid monolayer. Using this setup, we measured the motility of phospholipid molecules by performing Fluorescent Recovery After Photobleaching (FRAP) experiments for different lipidic compositions. The results of the FRAP measurements reveal that the motility of phospholipids is controlled by the monolayer packing decorating the interface.

## 1. Introduction

Lipid droplets are organelles present in the Endoplasmic Reticulum (ER) membrane [1, 2]. They attract many interest due to their role in several diseases like cancer and metabolic problems [3, 4, 5]. One of their functions is to store lipids and to regulate their consumption based on mechanisms that are not yet understood [6]. Lipid droplets (LDs) are composed of a core of neutral lipids and are surrounded by a phospholipid monolayer [7, 8]. Their biogenesis is assumed to occur in three consecutive steps [2, 9]. First, neutral lipids are synthesized in the ER membrane. Then, they accumulate and segregate into the ER bilayer core and form a lenticular droplet [2, 10, 11]. And finally, they grow through the accumulation of lipids until budding and sometimes pinch-off from the ER membrane [2, 12, 13, 14, 15]. Many of the biophysical properties of LDs have been investigated over the last decade using model systems [6]. As example, droplet interface bilayers (DiB), advanced free-standing bilayer (AFB), and giant unilamellar vesicles (GUVs) have been extensively used to reconstitute artificial and natural LDs in such lipid membranes [16, 17, 18, 19, 20, 21]. After successful reconstitution of LDs in a bilayer, the morphology of LDs was studied as function of the phospholipidic composition of the DiBs, GUVs or AFB systems [18, 19, 20]. It has been shown that the shape of LDs in a bilayer is determined by the balance of surface forces at the contact line between the embedded-LDs and the bilayer [18, 19, 20]. Thus, the associated contact angle of LDs can be predicted using wetting theory [11, 22]. Accordingly, numerical simulations based on continuum models were able to confirm many experimental results obtained on microscopic LDs [15, 18, 23, 24, 25, 26, 27].

In addition to the equilibrium shape determined by a force balance, nonequilibrium phenomena related to LDs have also attracted attention in recent years. A key example is protein partitioning, which is used to understand the dynamic distribution of specific proteins between the monolayer covering the LD and the bilayer in which the LDs are embedded [6, 16]. Indeed, protein partitioning seems highly dependent of the physical properties of the lipid monolayer (packing, fluidity) that surrounds the LD surface [18, 17]. However, to the best of our knowledge, there are no experimental measurements on phospholipid properties such as motility at the LD surface. One reason for this lack of knowledge could be the small size of the reconstituted LDs in the artificial systems (DiB, AFB, GUVs), which make FRAP experiments and extraction of precise diffusion coefficients challenging [17, 28]. As a consequence, there are only a few experimental studies in the literature that address diffusion. [24]. Interestingly, this fact is not limited to experimental studies but also occurs in theoretical studies with numerical simulations. [18, 25, 29].

In this article, we experimentally investigate the motility of phospholipids covering the interface between an aqueous and an oily phase. For this purpose, we have developed a microfluidic setup that enables the formation of a triolein-buffer interface decorated with phospholipids and have performed FRAP experiments directly at this interface (Fig. 1). Here, the planar phospholipid monolayer at the triolein-buffer interface mimicks the surface of a LD. We used triolein as neutral lipid because it is an important component of the biological LD [5] and varied the molecular composition of the monolayer to assess the influence of the most common molecules that make up the LD surface in living cells. These molecules are mainly phospholipids and cholesterol. Because native biological LDs may contain cholesterol ester [30, 31], we also investigated the influence of LD-core composition on phospholipid motility by adding some cholesterol ester (chol-ester) to the neutral lipid (triolein). Our measurements show that the motility of amphililic lipids is directly influenced by the packing of these molecules coating the interface.

**Figure 1:**
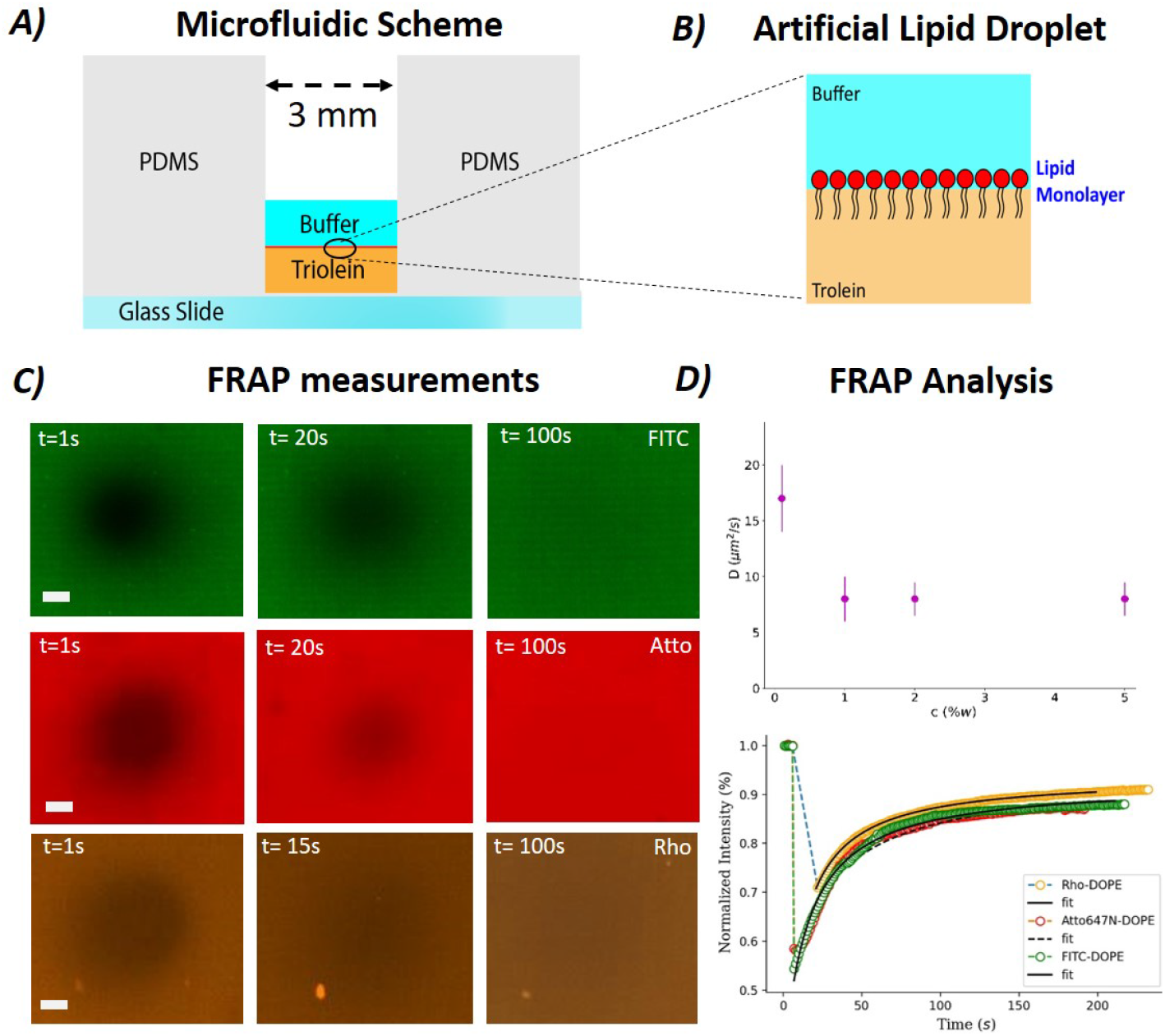
A) Sketch of the microfluidic device. B) Schematic of the phospholipid decorated triolein-buffer interface. C) Examples of FRAP measurements with FITC-DOPE (top), Atto647N-DOPE (middle) and Rho-DOPE (bottom). The time *t* = 0 corresponds to the moment when the bleaching was stopped. The length bar denotes 50 *μm* and corresponds to the radius of the bleached area. D) Upper panel: Measured diffusion constant for a pure DOPE monolayer with 2% Atto647N-DOPE (in molar ratio) for different DOPC concentration (expressed in weight ratio). Bottom panel: Experimental intensity data, as obtained from a FRAP measurement plotted together with the Soumpasis equation fitted to these data (dashed line); orange dots represent Rho-DOPE, red dots represent Atto647N-DOPE and orange dots represent FITC-DOPE [32]. Fits correspond to continuous and dashed dark lines (and all fits yield *D*≈17μm^2^*.m^-1^*).

## 2. Results and Discussion

Liquid droplets (LDs) originate in the ER membrane, and the phospholipid monolayer that decorates LDs is therefore composed mainly of phospholipids, sterols and proteins [33, 34, 35, 36, 37], with phospholipids making up the vast majority and proteins and sterols being present in only a small percentage. The decorating phospholipids can be distinguished in two families, saturated and unsaturated. Accordingly, we start our discussion with the most common lipids and presented the results sorted by lipid families: unsaturated phospholipids, saturated phospholipids, and sterols.

### Unsaturated Phospholipids

DOPC, DOPE, N-DOPE, POPC, LPC. A relevant biophysical composition for a phospholipid monolayer covering LDs localized in the ER membrane can be approximated by a DOPC/DOPE mixture with a molar ratio between 1:1 and 3:1 [7, 8, 17, 18]. For completeness, we vary the composition from pure DOPC to a pure DOPE monolayer. The extracted diffusion constants are presented in Fig. 2 and Table 1. The diffusion constant of a pure DOPC monolayer is *D* ≈ 8 *μm^2^* · s^-1^, i.e. close to, or slightly lower than, the DOPC fluidity observed in giant vesicles

**Figure 2:**
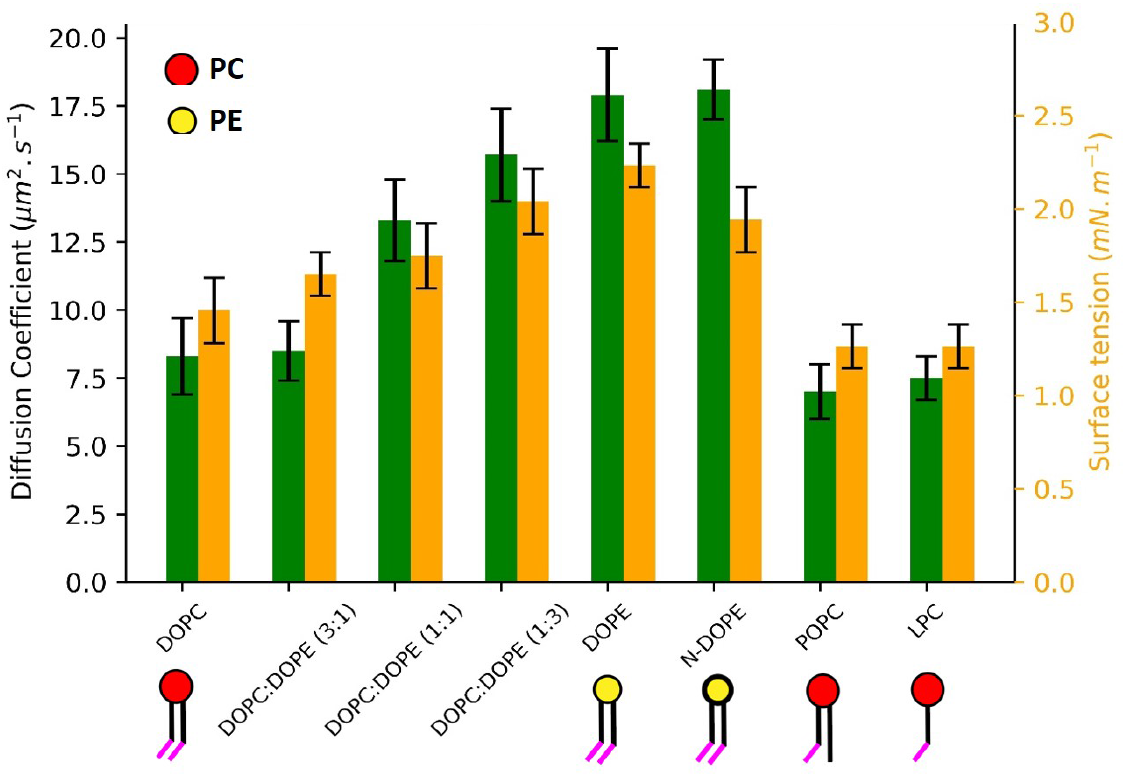
FRAP experiments yielded diffusion constants *D* and surface tension values for several saturated phospholipids (DOPC, DOPE, N-DOPE, POPC, LPC) and mixtures of DOPC and DOPE at a triolein-buffer interface. Schematic representations of the various lipids are provide for PE (red head), PC (yellow head) where pink tails indicate one unsaturation.

**Table 1:** Table summarizing the measured surface tensions *γ* and diffusion constants D for triolein-buffer interfaces decorated with monolayers made of a single lipidic component. More results including lipid mixtures are available in the plots shown in Figs. 2,3 & 4.

| Lipids | DOPC | DOPE | POPC | DPPC | DSPA | SM |
| --- | --- | --- | --- | --- | --- | --- |
| $\gamma$ (mN/m) | $1.5 \pm 0.3$ | $2.3 \pm 0.3$ | $1.3 \pm 0.3$ | $1.1 \pm 0.3$ | $2.6 \pm 0.3$ | $1.2 \pm 0.3$ |
| $D$ ($\mu\text{m}^2/\text{s}$) | $8.2 \pm 1$ | $17.1 \pm 1.6$ | $7 \pm 1$ | $6 \pm 0.6$ | $20.1 \pm 1.8$ | $8.1 \pm 0.9$ |

To also compare the observed motility of a DOPE monolayer with that of a bilayer, bilayer motility was determined using a free-standing DOPE bilayer as described in [28]. The formed free-standing DOPE/DOPC bilayers (with a 1:1 molar ratio) were formed using a 3D microfluidic chip as described in detail in reference [28]. Interestingly, the diffusion constant determined by FRAP measurements on a free-standing DOPE/DOPC (1:1) bilayer with 2% DOPE-Atto647 (in molar ratio) reveals ≈11 μm^2^/s, which is very close to the diffusion constant in giant vesicle (≈10μm^2^/s) but lower than that of a DOPE monolayer [38]. So in contrast to monolayer, no clear dependence on the bilayer motility was observed for different lipids.

The observed dependence of monolayer motility on the type of lipid can be easily understood considering the lipid packing that results from the geometry of these two molecules. DOPC can be approximated as a cylinder, while the shape of DOPE is closer to a truncated cone [39]. Assuming a cylindrical shape, the volume of a single phospholipid molecule can be approximated by the cross sectional area of its head *a*_0_ and its tail length *l*, as V_DOPC_ ≈ l · *a*_0_ for DOPC and 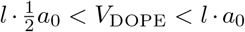 for DOPE. Thus, the corresponding critical packing parameter *ρ* = *V/a_0_ · l* is ρ_DOPC_ ≈ 1 for a monolayer consisting only of DOPC molecules and 0.5 < ρ_DOPE_ < 1 for a monolayer consisting only of DOPE molecules [39]. For a flat interface, a DOPC monolayer obviously has a higher lipid packing than a DOPE monolayer. In consequence, at equivalent surface coverage, the average motility of a single DOPE molecule is larger than that of a single DOPC molecule. Accordingly, packing density and mobility of DOPC and DOPE mixtures interpolate between the limits of a pure DOPC and DOPE monolayer, respectively.

As the increased lipids density at the triolein-buffer interface lowers the surface tension, differences in surface coverage can also be confirmed by differences in surface tension measured by the hanging drop method as γ _DOPC_≈1.5 mN/m and γ_DOPE_ ≈ 2.3 mN/m, for DOPC and DOPE, respectively. These surface tension values are consistent with expectations based on geometry dependent lipid packing and the experimentally observed lipid motility.

To confirm the influence of the lipid packing on lipid motility in a monolayer, we repeated the experiments described above with monolayers consisting of N-DOPE, POPC and LPC molecules. Compared to DOPE, has N-DOPE an additional methyl group in the PE head, thus this molecule is slightly less conical than DOPE but also less cylindrical than DOPC (Fig. 2). The measured diffusion constant for a pure N-DOPE monolayer is *D ≈* 17.5 μm^2^ · s^-1^ and the corresponding surface tension is ≈2 mN/m. As expected from the N-DOPE geometry, are both diffusion constant and surface tension rather comparable to pure DOPE monolayer.

POPC has the same hydroplilic head as DOPC and the same length of the hydrophobic tail, but only one unsaturation in its tail [34, 35]. Thus, due to this little difference, a POPC monolayer is expected to present a similar packing than a DOPC monolayer [27, 33, 40, 41]. And consistently with the geometry of the molecules and the expected monolayer packing, we measured the diffusion constant and the surface tension of POPC as *D* = 7 *μm^2^* · s^-1^ and γ ≈ 1.3 mN/m, that are very similar to those of a pure DOPC monolayer, see Fig. 2.

LPC is a single tail lipid with one unsaturation in its tail and an inverted conical geometry. The measured diffusion constant for a pure LPC monolayer is *D* ≈ 7 μm^2^ · s^-1^ and its surface tension is *γ ≈* 1.3 mN/m, see Fig. 2. Thus, this LPC appears to have a motility close to POPC and DOPC. This result is a little surprising, as *a priori*, we may have expected to measure a stronger difference between a monolayer composed of lipids with a single tail or a double tail. However, the correlation between low surface tension resembling high lipid packing and low lipid motility still holds true.

### Saturated phospholipids (DPPC, DSPA) and Sphingolipids (SM)

The motility of unsaturated lipids that make up the majority of the lipids covering LDs [42] was explored in the previous section. In this section, we continue our study by exploring the motility of saturated phospholipids on the surface of model lipid droplets. Due to the absence of unsaturations, the packing of unsaturated (neutral) phospholipids may be expected to be denser than that of unsaturated phospholipids. Following our findings for unsaturated lipids, the corresponding motility of saturated lipids, like DPPC, is generally expected to be lower than that of DOPC. However, the degree of saturation is not the only interaction that controls the monolayer packing. As example, electrostatic interactions can also strongly influence the packing of lipid monolayers. For this reason, we also ex-

**Figure 3:**
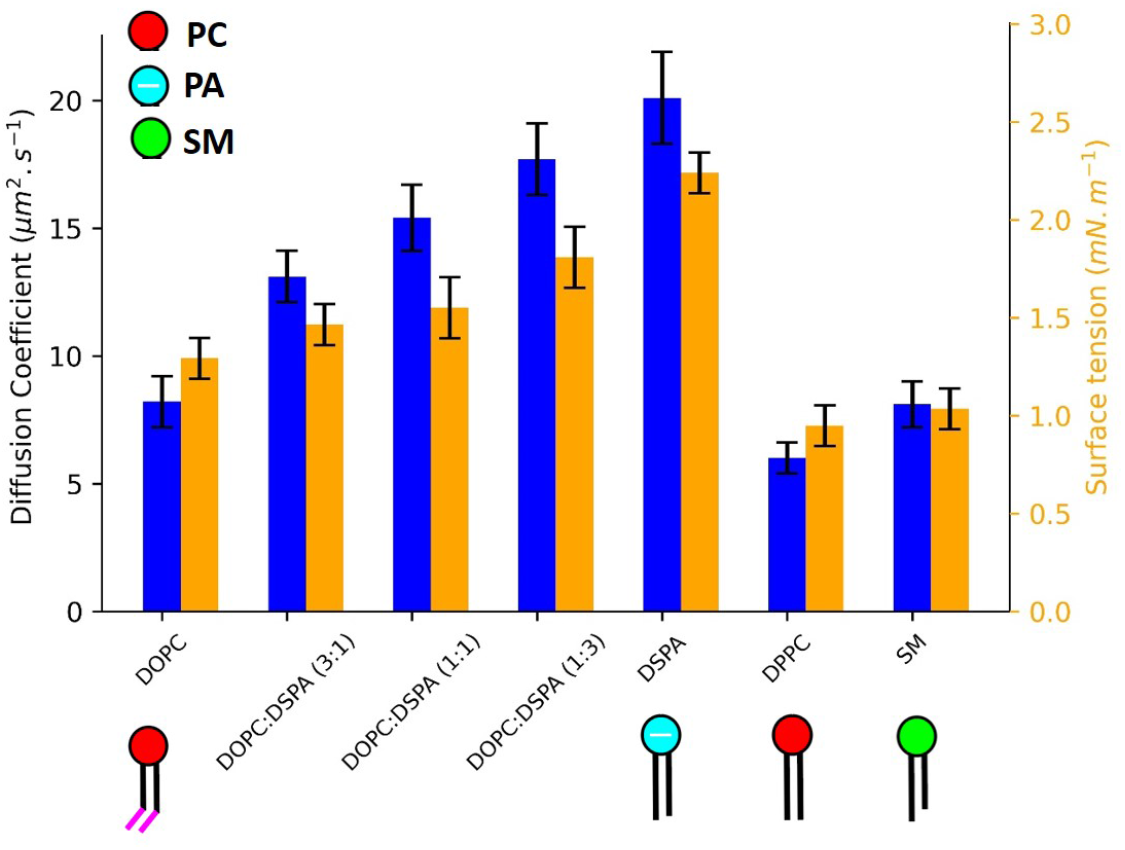
From FRAP experiments, diffusion constants *D* and surface tension values for unsaturated phospholipids (DSPA, DPPC, and SM) and mixtures of DOPC and DSPA were extracted and compared to DOPC values. A schematic representation is provided for PC lipids (red head), SM (green head) and the charged PA (light blue head).

plore mixtures between saturated DSPA molecules having an negatively charged hydrophilic head group and uncharged and unsaturated DOPC molecules. In addition to saturated lipids, we also consider the case of sphingolipids (SM).

DPPC is a saturated lipid with a cylindrical shape that self-assembles in an highly packed monolayer [43]. The measured diffusion constant for a pure DPPC monolayer is *D ≈* 5.5 μm^2^ · s^-1^, which is a little slower to that of unsaturated DOPC and POPC phospholipids, that are also expected to be densely packed (see, Fig. 3). The high DPPC packing at the triolein-buffer interface also coincides with a low surface tension, *γ ≈* 1.1 mN/m, which is also very similar to surface tension values for DOPC and POPC.

DSPA is a saturated double-tailed lipid with a head that is negatively charged with a conical shape. Due to electrostatic repulsion, the lipid packing of a pure PA monolayer is smaller than that of other lipids [44]. In addition to a large motility of *D ≈* 20 μm^2^.s^-1^ of a pure DSPA monolayer, the comparatively low packing density also leads to a comparatively large surface tension at the triolein-buffer interface, which was measured to be *γ ≈* 2.6 mN/m, see table 1. Mixtures of DOPC and DSPA molecules show motilities intermediate between those of a pure DOPC and a pure DSPA monolayer. Presumably due to the repulsion of the charged DSPA molecules, the measured diffusion constants are already quite large at a DSPA content of 25% and increase even further with increasing DSPA content, see Fig. 3.

Sphingolipids are a family of lipids that are synthesized in the (ER) membrane, where they can be found in low amounts, as well as at the trans Golgi [45,46, 47]. Due to the possible role of sphingolipids in the LD biogenesis, they have attracted a growing attention in recent years [45, 46, 47]. Sphingomyelin (SM) has a PC head with a fatty acid group and a tail that has a single saturation in contact with the head. Thus, we study this lipids with other saturated lipids. SM molecule has a slightly conical geometry close to a cylindrical molecule. The relatively low diffusion constant of a pure SM monolayer of *D ≈* 8 μm^2^ · s^-1^, and the corresponding low surface tension γ≈1.2 mN/m, Fig. 3, indicate that despite of the slightly conical shape of the SM, the associated lipid packing is dense, similar to an assembly of cylindrical molecules, in agreement with literature [39, 48].

### Sterol and Sterol ester

Sterols like cholesterol (chol) and cholesteryl ester (chol ester) are also important components of LDs [7, 8]. However, these molecules differ from previous lipids, as they are unable to form a lipid bilayer on their own. As example, chol ester is not a surface active molecule, it is a component of the LD core and not of the LD surface. Nevertheless, these sterols are important lipids and it is interesting to investigate their influence on the properties of phospolipid monoloayers..

Cholesterol (Chol) is present in the ER membrane only in small amounts, namely below 6 % percent. Nevertheless, Chol may play a role in metabolic diseases that are involving LDs [4, 49]. Chol drastically alters the biophysical

**Figure 4:**
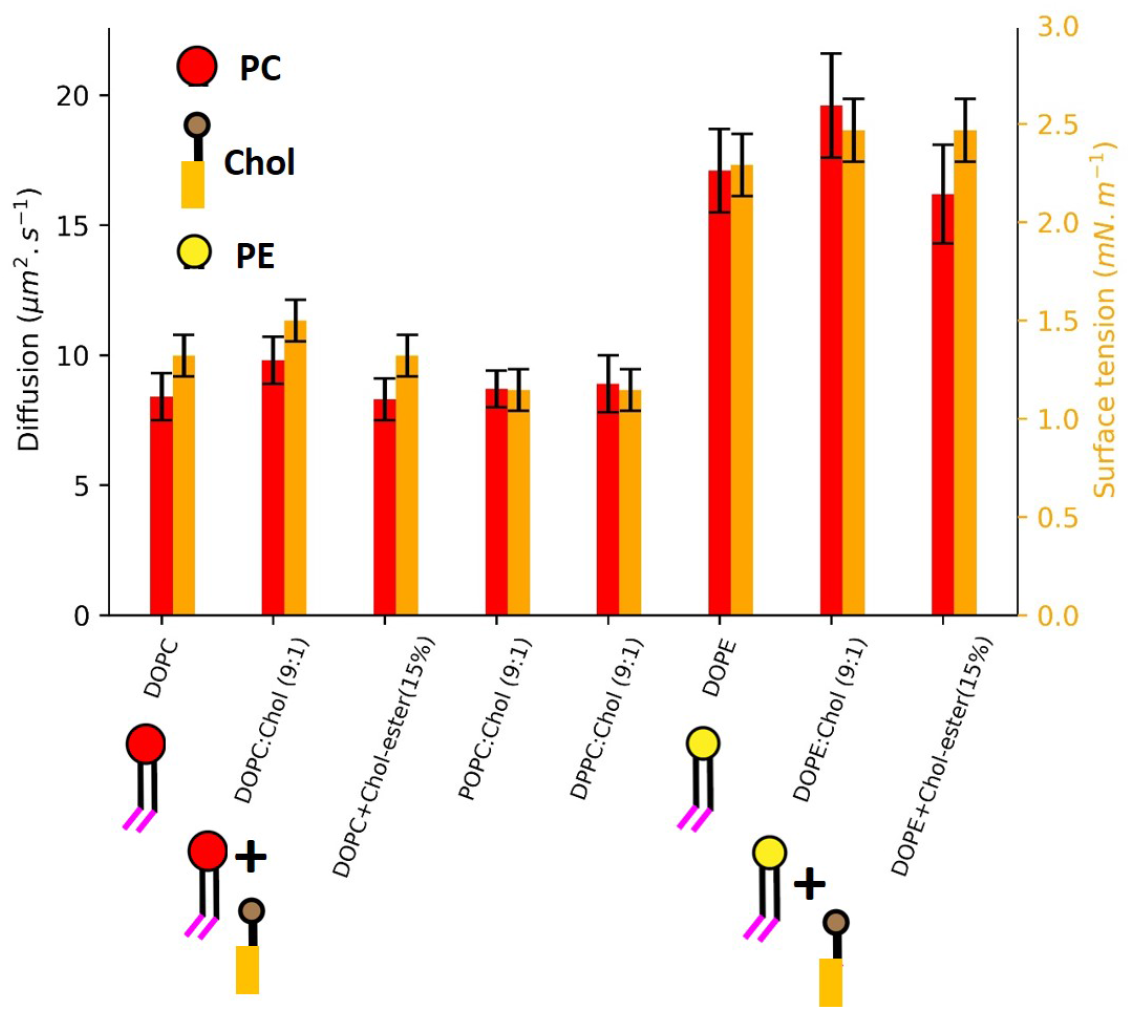
Diffusion constants D are derived from FRAP experiments and surface tension values for cholesterol-containing DOPC, DOPE, POPC, and DPPC monolayers at the triolein-buffer interface, as well as DOPC and DOPE monolayers at the cholesterol-ester interface. Schematic representation is provide for cholesterol and DOPC.

properties of the plasma membrane and thus, it is interesting to investigate the case of a lipid monolayer enriched with Chol. It is known that Chol increases both the bilayer tension and the bilayer fluidity, at least at low temperatures [50,51, 52]. The measured surface tension for a DOPC monolayer with 10 % Chol is ≈1.7 mN/m, while the surface tension of a DOPE monolayer with 10 % Chol is ≈2.5 mN/m, i.e. the surface tension for both phospholipids are increased by about 10 % compared to pure DOPC or DOPE monolayer, respectively, see table 1. The observed increase in surface tension indicates a reduced packing density of lipids, which is expected to increase lipid diffusion. In fact, the measured diffusion constant for a DOPC monolayer with 10% Chol gives a value of ≈10 *μm^2^·* s^-1^, while that of a DOPE monolayer with 10% Chol gives ≈20 μm^2^·s^-1^, see Fig. 4, i.e. the diffusion constant also increased by about 10 % compared to pure DOPC or DOPE monolayer. This suggests that the increase in surface tension and lipid diffusion is indeed due to decreased lipid packing.

Cholesteryl ester, a dietary lipid, is an ester of cholesterol found in LDs [53]. This molecule differs from the previous molecules in that it is not localized at the surface of LDs but it is only present in the LD core. This is interesting because cholesterol ester appears to affect the recruitment of some LD proteins when they enrich the LD composition [30, 31]. The surface tension of a DOPC monolayer at the interface between triolein-buffer with 15 % cholesteryl ester in triolein was measured to be γ≈1.5 mN/m, i.e. the same value as for triolein not containing cholesteryl ester. Similarly, we also measured the surface tension for a DOPE monolayer to be ≈2.3 mN/m no matter if the triolein contains 15 % cholesteryl ester or not. The corresponding measured diffusion constant yields a value of *D* ≈ 8 μm^2^·s^-1^ for a DOPC monolayer with 15 % cholesteryl ester in triolein, and a diffusion constant *D* ≈17 μm^2^·s^-1^ for DOPE monolayer with 15 % cholesteryl ester in triolein, see Fig. 4. From this, we can conclude that cholesteryl ester molecules remain in the triolein phase and hardly change the surface properties of the lipid decorated triolein-buffer interface. Similarly, we observed no measurable change in the surface tension of the phospholipid-decorated triolein-buffer interface or in the corresponding diffusion constant compared to a pure DOPC or a pure DOPE monolayer, respectively.(cf. Fig. 4).

## 3. Conclusion

We investigated the diffusion properties of lipids on the surface of model lipid droplets. For that, we produced a model lipid droplet interface made of a triolein-buffer interface in a microfluidic chamber that is decorated with a lipid monolayer of variable composition. FRAP experiments were conducted directly at this interface and the corresponding diffusion constants for the lipids were obtained as function of the molecular composition of the lipid monolayer. Based on geometrical (cylindrical, conical) and chemical (electrostatic charge, saturation) properties of the respective lipids, we argued that the lipid packing is the key parameter to understand the measured results. In this context, monolayers with high lipid packing have low lipid motility. And monolayers with low lipid packing have high lipid motility. This reasoning is supported by surface tension measurements as high lipid packing means low interfacial tension and low lipid packing means high interfacial tension. By extrapolation, this study provides an estimate of defect density on the LD surface for the lipids composition tested. This point is important because defects seem to play a key role in protein partitioning on LD surface [17].

## 4. Methods

*Molecules*. For the preparation of molecular monolayer that cover the model lipid droplets, i.e. the triolein-buffer interface, the following molecules were purchased from Avanti Polar Lipids: 1,2-dioleoyl-sn-glycero-3-phosphocholine (DOPC), 1,2-dioleoyl-sn-glycero-3-phosphoethanolamine (DOPE), 1,2-distearoyl-sn-glycero-3-phosphate (sodium salt) (DSPA), 1-palmitoyl-2-oleoyl-sn-glycero-3-phosphocholine (POPC), 1-(10Z-heptadecenoyl)-2-hydroxy-sn-glycero-3-phosphocholine (LPC), 1,2-dipalmitoyl-sn-glycero-3-phosphocholine (DPPC), 1,2-dioleoyl-sn-glycero-3-phosphoethanolamine-N-methyl (N-DOPE), cholesterol (Chol), N-stearoyl-D-erythro-sphingosylphosphorylcholine (SM), 1,2-dipalmitoyl-sn-glycero-3-phosphocholine (DPPC), Cholesteryl ester (Chol ester). Glyceryl trioleate (triolein) and squalene were purchased from Sigma-Aldrich. Recombinant human denaturtared perilipin 2 protein (PL2) was purchased from Abcam (ab181932). To allow for FRAP experiments, the following fluorescent labeled lipids were used: (FITC-DOPE) 1,2-dioleoyl-sn-glycerol-phosphoethanolamine Fluorescein purchased from ChemCruz (Santa Cruz Biotechnology), 1,2-dioleoyl-sn-glycero-3-phosphoethanolamine-N-(lissamine rhodamine B sulfonyl) (ammonium salt) (Rho-DOPE) purchased from Avanti Polar lipids, and 1,2-Dioleoyl-sn-glycero-3-phosphoethanolamine labeled with Atto 647N (Atto647N-DOPE) purchased from Sigma-Aldrich.

### Interfacial Tension Measurements

The interfacial tension of phospholipid decorated triolein-buffer interfaces was determined by the pendant drop method using a contact angle measurement device OCA 20 (DataPhysics, Germany). For this method, a transparent cuvette is filled with the lower density fluid, here the triolein-phospholipid mixture (*ρ_oil_* = 0.91 g/cm^3^) and a pendant drop containing a defined volume of the denser fluid, here the buffer solution (*ρ_buffer_* = 0.998 g/cm^3^), is produced from a hollow needle immersed in this liquid. While gravity drags the droplet down, buoyancy and surface forces keep the droplet in place. Based on a contour fit to the pendant droplet, the software SCA 20 extracts both size and shape of the droplet and automatically computes the corresponding interfacial tension. This procedure is most reliable when the droplet size is close to detachment from the needle, and we thus set the droplet volume to (2.0 ±0.5) μl.

### μChip Fabrication and Monolayer Formation

Using computer software (Auto-cad), we created a drawing in the form of a cylinder with 3 mm diameter, which was produced using a 3D printer. This cylinder was positioned in the center of a glass Petri dish. After setting up the mold, mixed and degassed PDMS (Sylgard 184 – Dow Corning) was poured onto the mold with a resulting layer thickness of about 3 mm so that the cylinder is not fully immersed in the PDMS. After curing the PDMS at 60 °C for 6-7 hours, the cylinder was removed and square pieces with the cylindrical hole in the center were cut from the PDMS and removed from the Petri dish. The final device was fabricated by attaching the square PDMS piece to a PDMS coated glass cover slip (Sylgard 184 spin coated for 5 min at 3000 rpm) using plasma bonding (Diener electronics). The thus fabricated device was placed on a hot plate at 60 °C for 1 hour to achieve higher bonding strength.

To produce the triolein-lipid mixture, a specific amount of lipid was dissolved in Ethanol. This solution was then dried under vacuum. Triolein was added to the dried components and mixed in an ultrasonic bath. The resulting phospholipid concentration in triolein is always equal to 2% w/w, which is above the critical concentration needed to form stable droplet interface bilayers in triacylglycerol oil [54, 17]. A 2% molar ratio of FITC-DOPE as a fluorescent probe. Such concentrations are typically employed for the production of stable artificial lipid droplets in literature [55, 18].

To form a monolayer, 6 μl of the as-prepared triolein-lipid mixture was first placed in the cylindrical opening of the PDMS micro chip, and then 20 μl of 150 M KCl buffer was added on top. Within seconds, a phospholipid monolayer forms at the triolein-buffer interface, with a composition similar to the lipid composition of the originally prepared triolein solution.

### FRAP Experiments

Confocal images were acquired with an inverted microscope (Nikon Ti-Eclipse) with an Intensilight Epi-fluorescence light source and a laser unit (LU-NV, Nikon). The confocal microscope was equipped with a Yokogawa spinning disk head (CSU-W1; Andor Technology) and a fluorescence recovery after photo-bleaching module (FRAPPA; Andor Technology). Confocal imaging was conducted using excitation wavelengths of 481 nm (for FITC-DOPE 495/515) and 561 nm (for Rho-DOPE 560/583, Atto647-DOPE 643/665). The used emission filters have the wavelengths/bandwidths of 525/30 nm, 607/36 nm and 685/40 nm, respectively. For the FRAP experiments, a pinhole size of 50 μm was used with 40× oil objective having a working distance of 220 micrometer. Prior to bleaching, a circular stimulation area with diameter ranging between 5 and 50 micrometer was selected inside the bilayer (to increase measurements quality). Prior to a FRAP measurement, fluorescence imaging was performed for 20 s for each individual experiment. Then, bleaching was performed by increasing the laser power on the stimulation area to the maximum laser power for 20 s (70 mW for 481 and 561 nm, 125 mW for 647 nm) including 10 loops that were repeated for three times. During the recovery, image acquisition was continued for at least 5 min to be able to observe all the changes after recovery. Then, the obtained fluorescence recovery (FRAP) data are directly fitted with the Soupmasis equation using Python. All the experiments were performed at room temperature of 23°*C*.

Interestingly, similar FRAP curves (and identical bleached area) and fits were obtained for three types of dyes on a pure DOPE monolayer: FITC-DOPE, Rho-DOPE and Atto647-DOPE (see Fig.1). A notable difference between these dyes is that Rho-DOPE appeared more difficult to bleach than Atto647N-DOPE or FITC-DOPE.

## 5. Data availability

The datasets generated during and/or analysed during the current study are available from the corresponding author on reasonable request.

## 6. Authorship contribution statement

S.A: performed the experiments and analyzed the data. J.-B.F. designed the experimental setup and the research. All the authors discussed the results and wrote the manuscript.

## 8. Acknowledgements

Financial support is acknowledged by the Deutsche Forschungsgemeinschaft (DFG), SFB1027 (project B4).

